# mclUMI: Markov clustering of unique molecular identifiers enables dynamic removal of PCR duplicates

**DOI:** 10.1101/2025.03.15.643033

**Authors:** Jianfeng Sun, Shuang Li, Stefan Canzar, Adam P. Cribbs

## Abstract

Molecular quantification in high-throughput sequencing experiments relies on accurate identification and removal of polymerase chain reaction (PCR) duplicates. The use of Unique Molecular Identifiers (UMIs) in sequencing protocols has become a standard approach for distinguishing molecular identities. However, PCR artefacts and sequencing errors in UMIs present a significant challenge for effective UMI collapsing and accurate molecular counting. Current computational strategies for UMI collapsing often exhibit limited flexibility, providing invariable deduplicated counts that inadequately adapt to varying experimental conditions. To address these limitations, we developed mclUMI, a tool employing the Markov clustering algorithm to accurately identify original UMIs and eliminate PCR duplicates. Unlike conventional methods, mclUMI automates the detection of independent communities within UMI graphs by dynamically fine-tuning inflation and expansion parameters, enabling context-dependent merging of UMIs based on their connectivity patterns. Through *in silico* experiments, we demonstrate that mclUMI generates dynamically adaptable deduplication outcomes tailored to diverse experimental scenarios, particularly best-performing under high sequencing error rates. By integrating connectivity-driven clustering, mclUMI enhances the accuracy of molecular counting in noisy sequencing environments, addressing the rigidity of current UMI deduplication frameworks.

## Introduction

Genetic material is either fragmented for short-read sequencing or retained intact for long-read sequencing [1], laying the foundation for understanding genomic and transcriptomic expression levels across various experimental conditions [2]. Accurate quantification of these fragments or entire molecules is a prerequisite for better dissecting their expression profiles [3,4]. However, directly quantifying original molecules from sequenced reads is challenging due to the presence of polymerase chain reaction (PCR) duplicates [5,6]. Therefore, the identification and removal of PCR duplicates are crucial for accurately estimating the number of unique molecules [7].

Unique molecular identifiers (UMIs) are sequences of oligonucleotides that are randomly synthesised to tag unique molecules, enabling accurate molecular counting [8–11]. The incorporation of UMIs has become a reliable method of choice due to its proven feasible across various high-throughput sequencing experiments and its effectiveness in demultiplexing during downstream analysis [12–14]. PCR duplicates with the same UMI are assumed to originate from the same molecule, a process known as UMI collapsing or grouping, which facilitates the counting of unique molecules. However, errors are frequently introduced into UMI regions of both original molecules and PCR duplicates. This results in significant numbers of unique molecules tagged with different UMIs, despite sharing the same origin, complicating the UMI collapsing process. To address this issue, different UMIs are collapsed together within an edit distance to tolerate errors and ensure accurate identification of true molecules. This error-tolerant strategy is effective in grouping different UMIs when reads undergo PCR amplification and sequencing at low error rates. In several sequencing technologies, such as single-cell RNA-seq (scRNA-seq), reads are subjected to numerous PCR amplification cycles to obtain sufficient genetic material for sequencing. However, research has shown that the accuracy of calculating gene expression profiles declines significantly at high PCR cycles because the number of PCR artefacts [15]. Consequently, the accuracy of molecular quantification varies in sequencing conditions, highlighting the need for computational strategies in UMI collapsing that are dynamically adaptable to specific experimental conditions.

In recent years, numerous computational tools for UMI collapsing have emerged [16–23]. These tools primarily focus on removing PCR duplicates by constructing graphs of UMIs [24,25]. One of the most widely used tools, UMI-tools, exemplifies graph-based strategies for effectively identifying unique UMI-tagged molecules. The process begins by constructing a graph where each node represents a unique UMI. An edge is drawn between the two nodes if their edit distance is no greater than a specified threshold, *k*; otherwise, they remain unconnected. This approach generates multiple connected components within the graph, where the number of components roughly corresponds to the unique UMI count. This is known as the *Cluster* method in UMI-tools. Within each connected component, further subdivision can occur. The number of subcomponents, where each central node (having the highest count) is one edge away from the other nodes, is used to estimate the deduplicated molecule count. This approach is referred to as the *adjacency* method in UMI-tools. Since *cluster* and *adjacency* often led to an underestimated number and an overestimated number of unique UMIs, the *Directional* method was developed to seek for a more balanced estimation between somewhat of the two extremes. Its deduplication process can coarsely be described as a directed edge-visiting strategy, that is, merging node B by node A if the count of node A is at least two-fold greater than that of node B. The *directional* method in the *UMI-tools* suite is reported to gain the highest accuracy in identifying PCR duplicates [24]. However, regardless of the error settings or conditions, these strategies collapse UMIs into a single outcome once a set of reads is provided, potentially lacking the flexibility to dynamically expand or contract unique UMI counts to adapt to varying conditions.

Here, we present mclUMI, a graph clustering-based computational tool built using Markov clustering (MCL) algorithms [26–28], for precise localisation of unique UMIs and removal of PCR duplicates. Unlike established methods, the MCL-based approach identifies independent communities of UMIs within a connected component, each with its UMI nodes strongly connected and highly relevant. This process allows UMI nodes to be merged dynamically based on the connectivity of edges within a UMI graph. To incorporate the influence of UMI counts on the effectiveness of UMI collapsing, we developed two variant strategies, MCL-ed and MCL-val. Through comprehensive evaluations on simulated datasets with ground-truth information, we demonstrate that mclUMI achieves superior accuracy in molecular quantification compared to existing graph-based methods. As a complementary tool to state-of-the-art UMI identification frameworks, mclUMI enhances their applicability across heterogeneous experimental contexts.

## Results

### Markov clustering for UMI collapsing and PCR artefact removal

We developed mclUMI for automatic detection of UMI clusters by the Markov clustering algorithm without the need for calculating UMI counts, leading to the MCL method. It has two derivatives (MCL-ed and MCL-val) by considering the information about UMI counts (**Figure 1a**). As seen in **Figure 1b**, PCR artefacts or duplicates result from amplification of reads over multiple PCR cycles. Classification of PCR duplicates according to the same UMIs does not work when UMIs are synthesised and/or sequenced incorrectly because of DNA polymerase/sequencing errors. To tackle this problem, graph-based computational strategies have previously been proposed that leverage edit distances between UMIs to build UMI graphs (see [24] for technical review). Differently, mclUMI implements a Markov clustering algorithm for UMI collapsing based on a UMI graph. We began by simulating 7 UMIs for 10X V2 chemistry and constructing a UMI graph using an edit distance 1. Upon receiving the adjacency matrix of the UMI graph, mclUMI is asked to search for subcomponents with their respective nodes strongly connected (**Figure 1c**). For instance, a UMI graph composed of 7 nodes is further split into two subcomponents (**Figure 1d**). It shows high connectivity within each of them but low connectivity between them. Nevertheless, Markov clusters are discovered without leveraging the information on UMI counts. To include this information in MCL-based UMI collapsing, we additionally designed MCL-val, whose main thrust bears a very close resemblance to the Directional method. Rather than collapsing UMIs at the node level, MCL-val collapses UMIs at the subcomponent level. MCL-val identifies UMIs with the highest count from Markov clusters and merges two Markov clusters if the count of one of the most frequently observed UMIs is at least twofold higher than that of the other one. As illustrated in **Figure 1d**, because the difference in counts between UMIs A and B does not meet this requirement, the two Markov clusters remain separated. Building on the subcomponent level, we developed MCL-ed, which merges two Markov clusters if the two most frequently observed UMIs are within a *k* edit distance of each other. For the 7-node UMI graph, the two Markov clusters are merged because UMIs A and B are within one edit distance.

**Figure 1.**
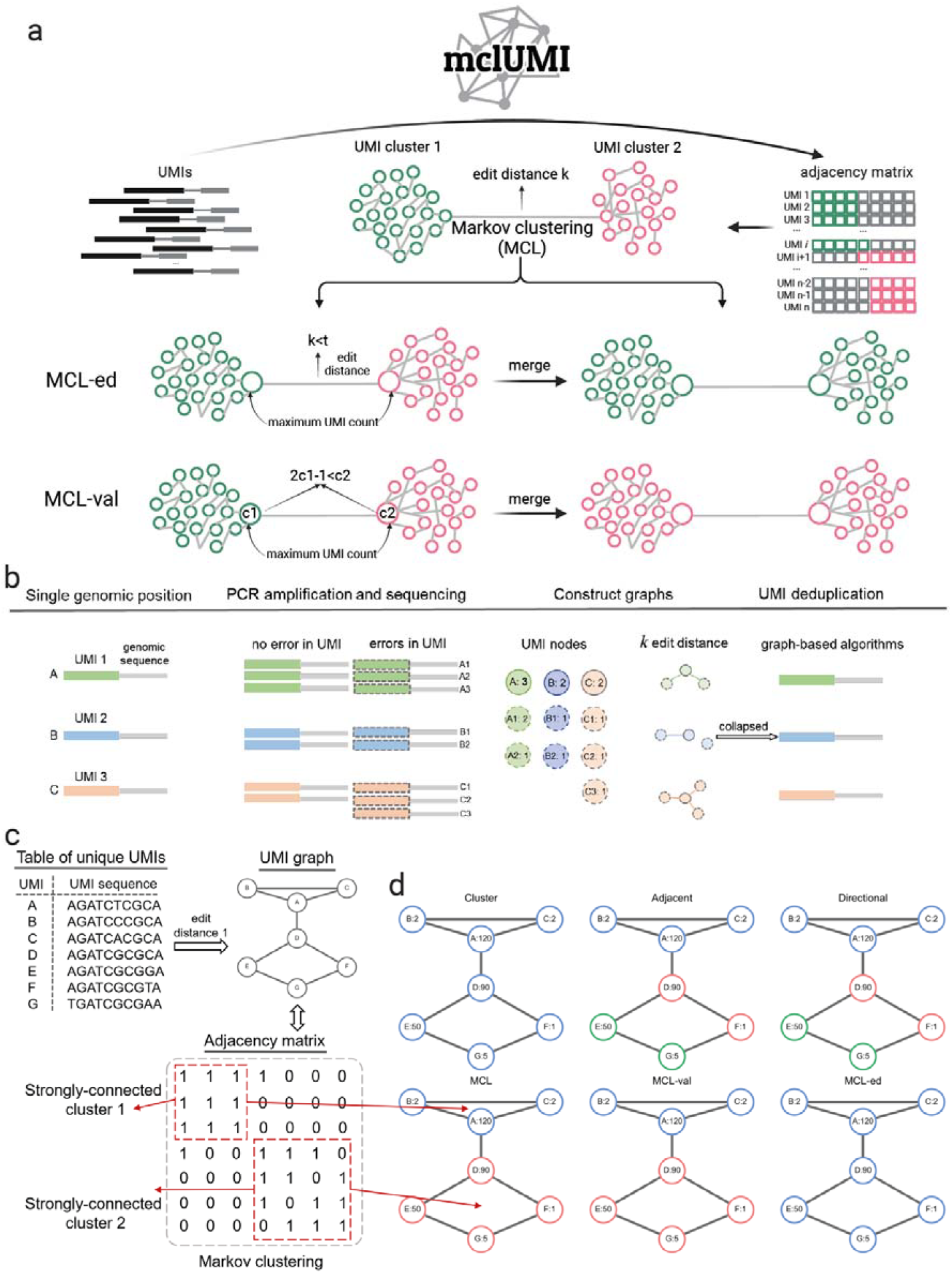
Overview of UMI deduplication using Markov clustering. **a**, Schematic of Markov clustering of UMIs in the mclUMI tool. **b**, Subjecting UMI-tagged molecules to PCR amplification and sequencing. The effectiveness of UMI deduplication is evaluated by how the estimated number of molecules is close to the actual number of molecules. **c**, Building a graph with 7 unique UMI nodes (A-G) connected with an edit-distance threshold of 1. Strongly connected components are visually displayed in the adjacency matrix. **d**, Graph-based identification of true molecules by collapsing UMIs using the cluster, adjacency, directional, mcl, mcl-ed, and mcl-val algorithms. PCR duplicates are merged into different UMI groups in different edge colours. The identifier of each unique UMI is followed by its count in the single genomic locus.

### Comparison of UMI collapsing performance

We next evaluated the performance of UMI collapsing methods on simulated reads under varying conditions, including PCR cycle numbers, PCR errors, sequencing errors, UMI lengths, PCR amplification rates, and sequencing depths. In addition to the graph-based clustering methods in mclUMI, we implemented several Euclidean distance-based clustering methods, including DBSCAN, Birch, and Affinity propagation. Unlike mclUMI, which uses an adjacency adjacency matrix as input (**Figure 2a**), the distance-based clustering methods rely on non-graphical data for clustering UMIs. As illustrated in **Figure 2b**, UMI sequences are represented by one-hot encodings, which are then flattened and fed into the distance-based clustering algorithms. Our results demonstrate that methods implemented in mclUMI, along with DBSCAN, Birch, and *Directional* show a clear advantage in UMI collapsing over *Unique* and *Adjacency* across all six sequencing settings. Notably, mclUMI provides better UMI collapsing results than *Directional* at high PCR and sequencing error rates (**Figure 2c**). Additionally, we found that without building bespoke UMI graphs, DBSCAN and Birch can achieve a deduplication profile similar to UMI-tools methods. Consistent with the findings of the UMI-tools study [24], we observed that the number of unique molecules deduplicated by is underestimated at certain thresholds, particularly concerning varying UMI lengths and sequencing depths.

**Figure 2.**
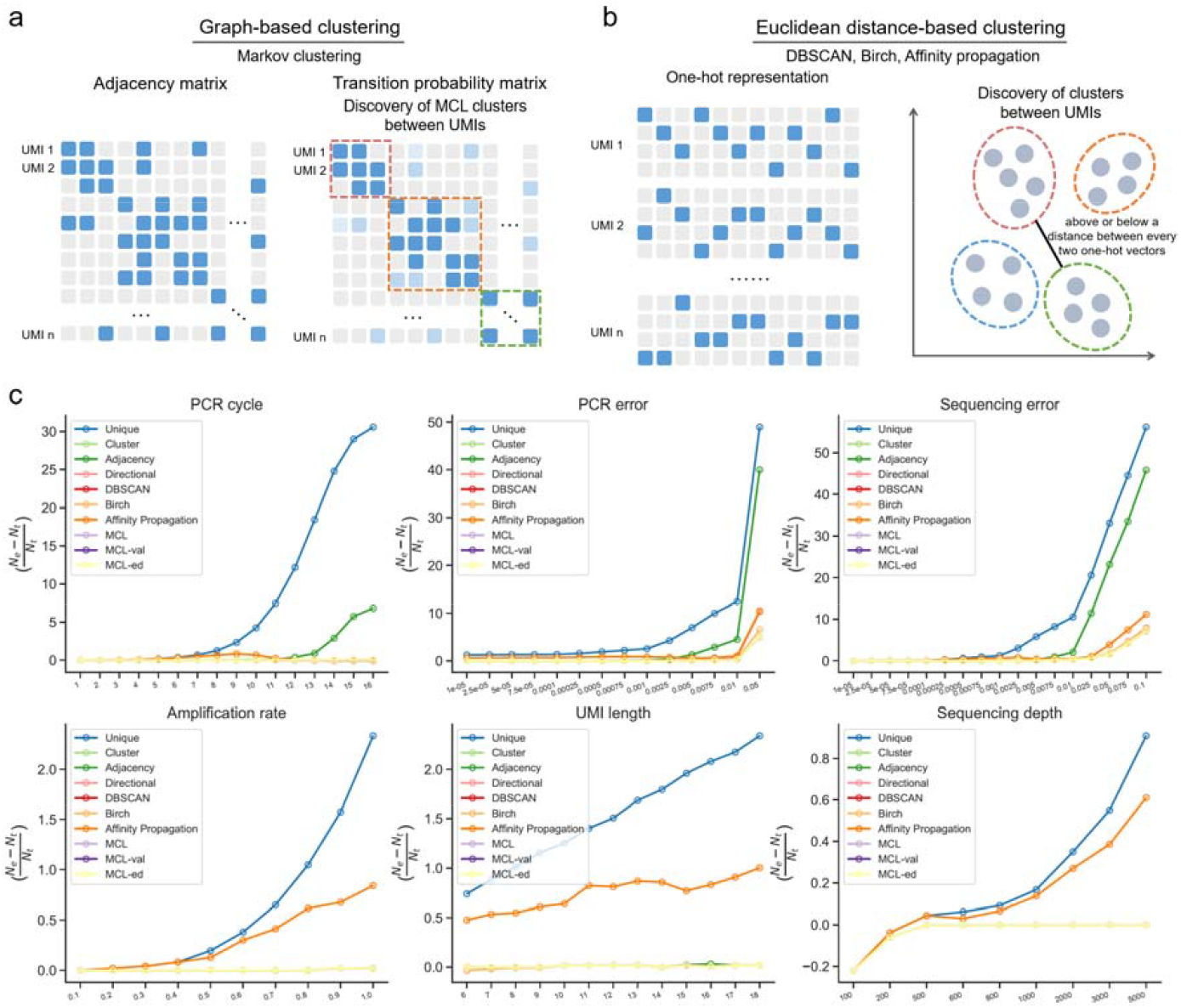
Performance comparison of UMI collapsing methods. **a**, Schematic of graph-based clustering methods for identifying UMI clusters based on the adjacency matrix and the transition probability matrix. **b**, Schematic of Euclidean distance-based clustering methods for identifying UMI clusters using one-hot representations of UMIs. One position in each column is occupied to be indicative of its nucleotide type. The one-hot matrix is flattened as a vector with a dimension of 1×4n (*n* represents the UMI length) prior to taking them as input to the Euclidean distance-based clustering methods. **c**, Comparison of deduplication results between methods across multiple PCR cycles, PCR error rates, sequencing error rates, amplification rates, UMI lengths, and sequencing depths, respectively. A close-up view of the deduplicated counts of Directional, MCL, MCL-ed, MCL-val, DBSCAN, and Birch can be found in **Supplementary Fig. 1**.

Additionally, we evaluated per-permutation UMI counts deduplicated by MCL-ed, MCL-val, and the UMI-tools *Directional* method across multiple sequencing error rates (**Supplementary Figure 2**). Our results show that both MCL-ed and MCL-val consistently yield lower fold changes in deduplicated UMI counts than the UMI-tools *Directional* method, highlighting that Markov clusters of UMIs can be post-processed and merged to further improve molecular quantification.

### Markov clusters of UMIs efficiently capture UMIs from the same origin

The UMI-tools *directional* method excels in distinguishing original UMIs from amplified erroneous, amplified UMIs, making it a benchmark for UMI collapsing by leveraging information about both edit distance and UMI counts. To merge UMI nodes, the method initiates a series of directed visits from the UMI node with the highest count to those with lower counts. Ideally, when a node merges a neighbouring node into a cluster, reads attached with the two UMIs are derived from the same original molecule. To assess the effectiveness of the *directional* method in merging UMIs from the same origin, we labelled each amplified read the identifier of its original molecule during the read simulation process. This allows us to track the source of each read. As shown in **Figure 3a**, there are four possible scenarios, UMIs that are merged and derived from the same origin, UMIs that are merged but derived from a different origin, UMIs that are not merged but derived from the same origin, and UMIs that are not merged and derived from a different origin. The ideal outcome is to merge two UMIs that share the same root. Our results show that most of the methods effectively collapse UMIs from the same origin with regard to merging nodes (**Figure 3b**). Since only the MCL-val, MCL-ed, and *Directional* methods involve custom-built conditions for determining whether to merge UMIs, we further calculated the percentage and count of UMIs that are not merged from the same or different origin. Interestingly, there is a significant increase in the number of UMIs not merged by MCL-ed when they are from different origins, whereas the *Directional* method exhibits the opposite trend (**Figure 3c-e**). In summary, these findings suggests that merging UMIs nodes at the subcomponent level helps avoid the inclusion of reads from different original molecules compared to merging at the individual node level.

**Figure 3.**
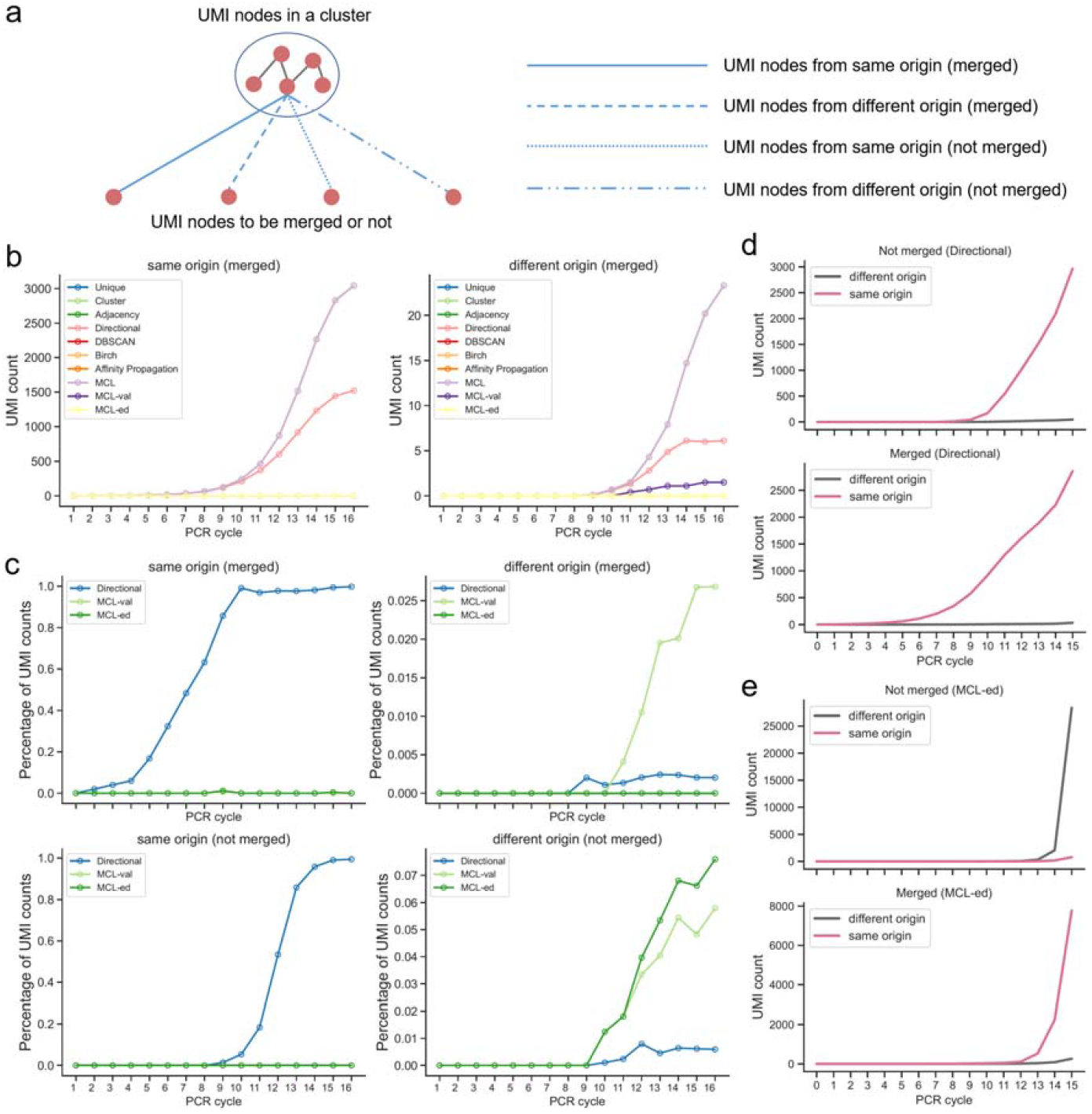
Researching trajectories of UMIs across PCR amplification cycles. **a**, Schematic of merging UMIs into a cluster. Annotating UMIs with their origin is conducive to study their connectivity in a network in which a merge indicates an estimated association between two UMIs. **b**, Counts of UMIs derived from the same origin to be merged across multiple PCR cycles with different UMI collapsing methods. **c**, Comparison of counts of merged or not merged UMIs derived from the same or different origin between the MCL-val/MCL-ed and Directional methods. **d** and **e**, Counts of merged or not merged UMIs derived from the same or different origin using the Directional (**d**) and MCL-ed (**e**) methods.

### Determination of optimal parameters of Markov clustering algorithms for UMI collapsing

The MCL algorithm utilizes two parameters, *inflation* and *expansion*, to discover various clusters from a UMI graph. To determine the best choice of the parameters, we varied *inflation* in a range between 1 and 6 and *expansion* in a range between 2 and 9 and deduplicated PCR artefacts on simulated reads across 6 sequencing settings. We found that both *inflation* and *expansion* induce the biggest perturbation on the deduplication effect when reads are subjected into a high PCR cycle number and large PCR/sequencing errors (**Supplementary Figure 3**). For dynamical clustering of UMIs, the *inflation* and *expansion* parameter are suggested to opt for a range between 1 to 3.5 and three integers 2, 3 and 4, respectively. Rather than an invariable result deduplicated by existing methods, this allows for generating various UMI clusters to achieve dynamic clustering.

### Molecular quantification on simulated scRNA-seq data

We subsequently compared the accuracy of molecular quantification between mclUMI and UMI-tools at the single-cell sequencing level. We first used SPsimSeq [29] to generate a single-cell count matrix composed of 10 cells and 10 genes under various sequencing error rates and then simulated sequencing reads only for genes expressed in cells using ResimPy [15]. We found that in approximately 67.5% of cell-by-gene positions, mclUMI provided better UMI collapsing results than the UMI-tools *Directional* method at sequencing error rates ranging from 0.02 to 0.05 (**Figure 4**). Then, to obtain a count matrix with known cell types for visualisation purposes, we used DeepConvCVAE—trained on the PBMC68k dataset [30] and hosted in UMIche (https://2003100127.github.io/umiche)— to simulate a 1100×20736 cell-by-gene count matrix where 100 cells for each of 11 known cell types were sampled (**Figure 5a**). We found that the single-cell map for both mclUMI and *Directional* methods are in high agreement with the reference map classified by t-distributed stochastic neighbour embedding (TSNE) [31], particularly, at high sequencing error rates 0.05 and 0.1 (**Figure 5b-c**).

**Figure 4.**
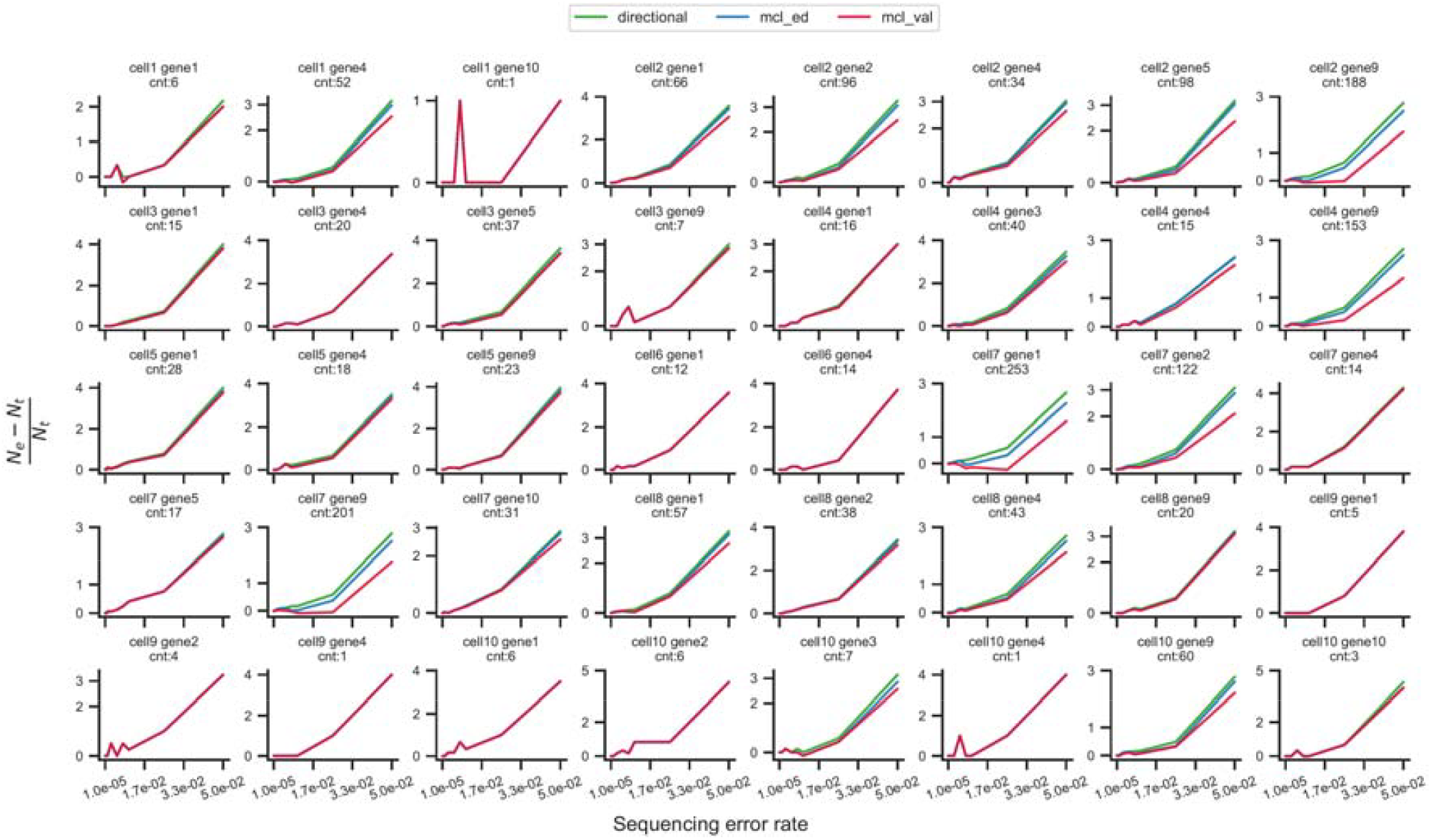
Comparison of UMI counting performance between mclUMI (MCL-ed and MCL-val) and the UMI-tools *Directional* method on single-cell sequencing data simulated by Resimpy at multiple sequencing error rates. Each subplot shows the UMI collapsing result for a certain cell and a gene. cnt: the count of a gene expressed at a cell.

**Figure 5.**
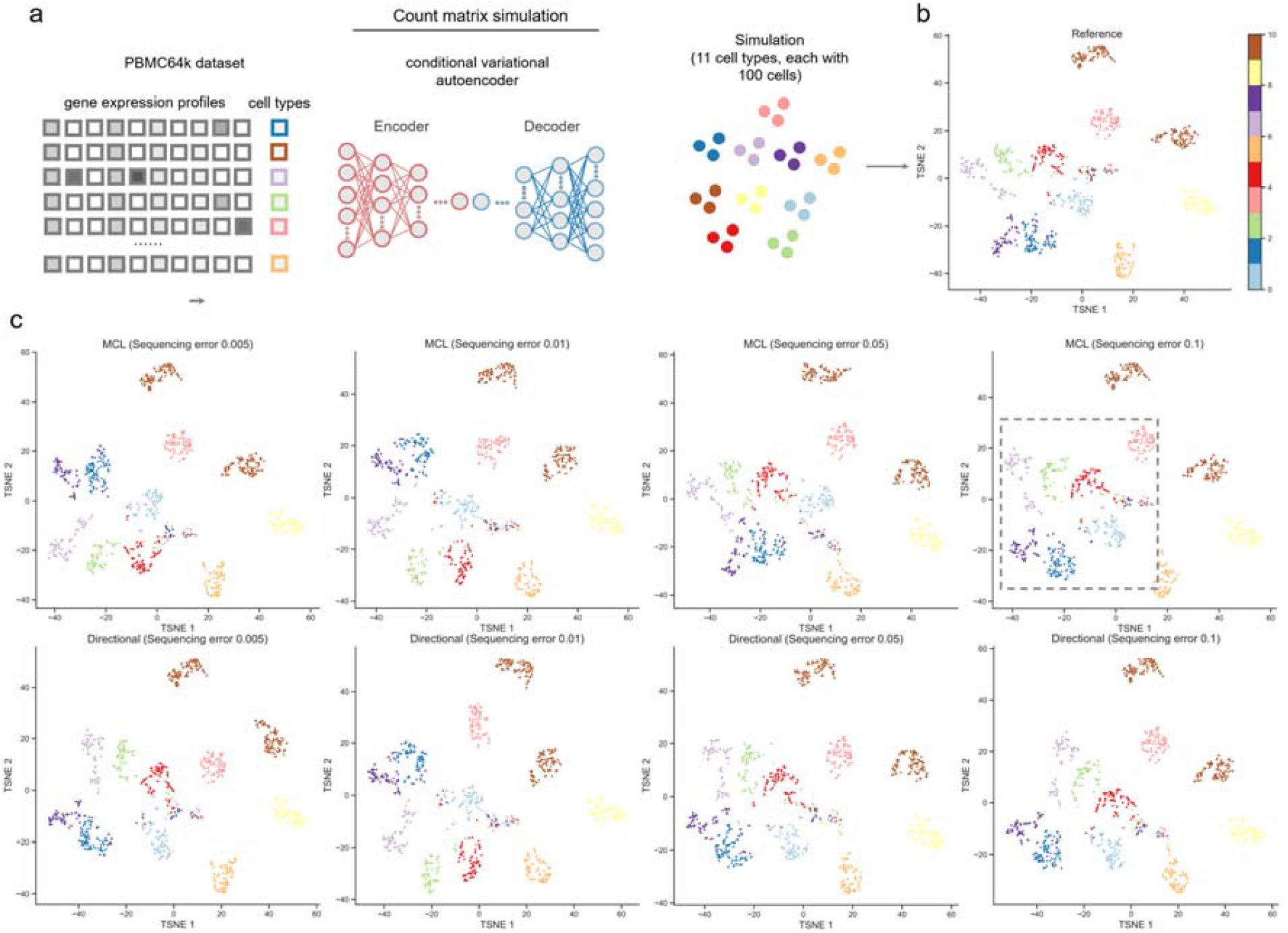
Clustering of deduplicated UMIs with single-cell sequencing data. A count matrix containing 100 cells for each of the 11 cell types is generated using our in-house count matrix simulator based on autoencoders. The resulting count matrix is then visualised by the TSNE method as reference. To estimate the counts per cell-by-gene type by UMI collapsing methods, we simulated the reads according to the count matrix through PCR amplification and sequencing. Subsequently, the sequenced reads with ground truth are respectively deduplicated by the Directional and MCL methods at 0.005, 0.01, 0.05, and 0.1 sequencing error rates. Ultimately, deduplicated counts for each cell-by-gene type are projected by TSNE.

### Case studies: connectivity of Markov clusters using mclUMI

Unlike existing methods, mclUMI emphasises the importance of identifying strongly connected subcomponents from a connected component. To illustrate how the connectivity between UMI nodes is determined, we analysed two connected components from simulated and iCLIP experiments [32], respectively. From simulated reads sequenced with an error rate of 0.05, we randomly selected a connected component with 67 unique UMIs a single edit distance apart. Using MCL, mclUMI gives two distinct subclusters, within which nodes are strongly connected but between which only two nodes are interconnected. By contrast, using the *Directional* method, UMI nodes are merged without considering the influence of the connectivity between UMIs, as indicated in the grey ellipse (**Figure 6a**). Similarly, **Figure 6b** illustrates that Markov clustering of 16 unique UMIs observed at a genomic position from the iCLIP dataset also captures strong connectivity between UMIs nodes. By contrast, the *Directional* method results in a few stray UMIs falling between or within clusters (e.g., the green UMI node). As UMIs in a cluster may be derived from the same UMI with the highest count, we projected UMI counts onto UMI nodes to understand whether Markov clusters capture not only strong connectivity but also UMIs which are assumed to be representative of clusters. Our *in silico* results agree to this assumption that there are two UMIs that are overrepresented in two major clusters in red and sienna, respectively.

**Figure 6.**
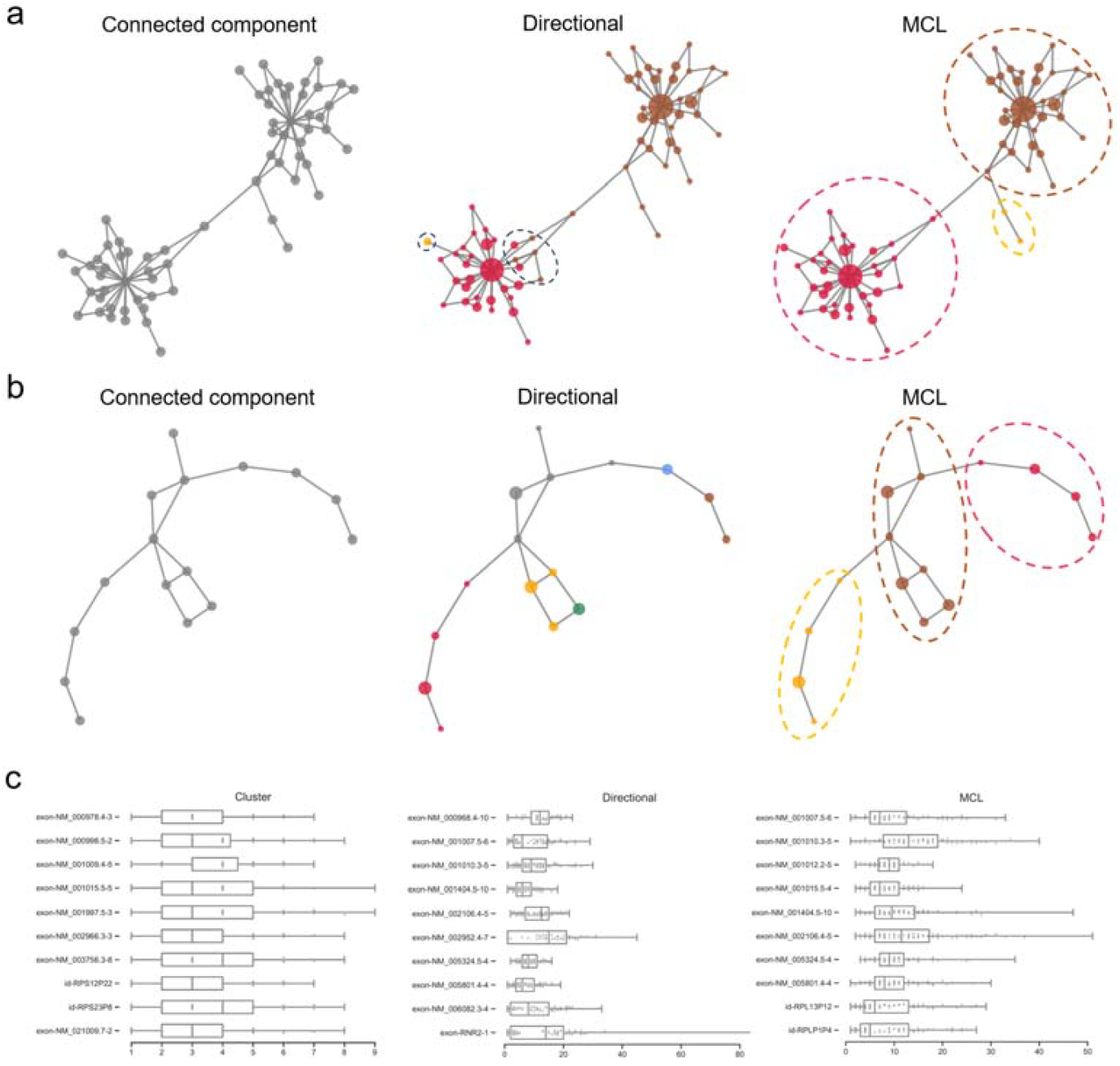
Quality evaluation and biological application of deduplicated UMI counts. **a** and **b**, UMI clusters observed at two example genomic loci using simulated data (**a**) and iCLIP data (**b**) [32], respectively, are discovered from connected components by the Directional and MCL methods. It contained 1,175,027 reads with 20,683 raw unique UMI sequences and 12,047 genomic positions tagged by running the get_bundles method of UMI-tools. Each node in the graphs represents a unique UMI. The node size is enlarged as the count of the unique UMIs increases. Different clusters are shown in different colours. **c**, Comparison of top 10 highly expressed genes identified by the Cluster, Directional, and MCL methods using a 10X genomics scRNA-seq dataset [33]. It contained 3,553,230 raw reads and left 588,963 reads following STAR (version: v2.7.9a) [46] mapping against GRCh38 and gene annotations using featureCounts (version: v2.0.1) [47]. The 100 barcodes were generated using the whitelist function of UMI-tools.

Furthermore, to assess how different UMI collapsing algorithms impact the identification of highly expressed genes, we plotted the expression profiles of the top 10 highly expressed genes using a single-cell dataset used in the original UMI-tools paper [24,33] (**Figure 6c**). We found that 6 out of 10 genes are identified identically by mclUMI and the *Directional* method.

## Discussion

We developed mclUMI, a Markov clustering-based toolkit to efficiently remove PCR duplicates. The method leverages two key parameters—inflation and expansion—which control the detection of Markov clusters with strongly connected UMI relationships, offering greater flexibility and adaptability compared to established methods. An original molecule is often PCR-amplified multiple times, resulting in a high-count UMI. We hypothesise that representative UMIs with the high counts across different clusters may share the same origin. This is because an erroneous UMI, introduced through DNA polymerase errors in early PCR cycles, can be preferentially amplified in later cycles, leading to its overrepresentation. To address this, we implemented two post-processing strategies:

MCL-ed (edit distance-based merging): If representative UMIs from different Markov clusters are within a minimal edit distance (*k*), they are connected via edges and merged based on sequence similarity.

MCL-val (Count-Based Merging): If representative UMIs from different Markov clusters have a count difference within a *t*-fold threshold, they are linked and merged based on UMI abundance.

Analysis of original molecule identifiers across multiple PCR cycles confirms that both strategies effectively distinguish true molecules from PCR duplicates, improving the accuracy of molecular quantification.

One limitation of mclUMI is its higher computational cost compared to the cluster, adjacency, and directional methods, as it requires iterative optimisation to refine Markov clusters. Additionally, our exploration of clustering-based UMI collapsing has led to the development of Euclidean distance-based clustering algorithms, with DBSCAN and BIRCH demonstrating strong performance in UMI deduplication.

The impact of sequencing errors is particularly pronounced in long-read sequencing technologies, where error profiles are magnified. Our results show that mclUMI outperforms UMI-tools at high sequencing error rates, providing a valuable complement to existing computational methods for PCR artefact deduplication.

## Methods

### Reimplementation of UMI-tools deduplication methods

The *UMI-tools* methods are most widely used in UMI identification. To better understand and compare graph-based UMI collapsing algorithms, we reimplemented all deduplication methods reported in *UMI-tools*, including *cluster, adjacency*, and *directional*. The *cluster* and *adjacency* methods were constructed using the breadth first search (BFS) algorithm while the *directional* method was constructed using the depth first search (DFS) algorithm.

The reimplementation process begins by constructing a graph where each node represents a unique UMI. An edge is drawn between the two nodes if their edit distance is no greater than a specified threshold,*k*; otherwise, they remain unconnected. This approach generates multiple connected components within the graph, where the number of components roughly corresponds to the unique UMI count. This is known as the *Cluster* method in UMI-tools. Within each connected component, further subdivision can occur. The number of subcomponents, where each central node (having the highest count) is one edge away from the other nodes, is used to estimate the deduplicated molecular count. This approach is referred to as the *Adjacency* method in UMI-tools. Starting from node *M* with the highest UMI count in each connected component, a neighbour *N* connecting the node is grouped into the same connected subcomponent using the UMI-tools *Directional* strategy if *C*_*M*_ is at least twofold larger than *C*_*N*_ where *C*_*M*_ and *C*_*N*_ are the counts of nodes *M* and *N*. Next, after the first subcomponent in the connected component is formed, the node with the highest count from all remaining unvisited nodes is picked up for forming the second subcomponent. This process is repeated and aborted until all nodes in the connected component are visited only once, leaving *K* subcomponents.

### Markov clustering of UMIs for collapsing

By calculating edit distances between UMIs and setting an editing distance cutoff, a graph is built to graphically present intricate relationships between UMIs. Edges that are used to link UMIs together reflect connectivity between them. If considering UMI counts as node sizes, such a graph can essentially reflect how UMIs with a low count evolve from a UMI with a high count over error-prone PCR amplification and sequencing processes. However, it remains murky as to which UMIs in a graph are most appropriately grouped and considered to originate from the same origin. To cope with this challenge, UMI-tools meticulously designed a set of custom-built, expert-guided rules to construct two methods, *Adjacency* and *Directional*. Different from this notion, we conjecture that edges are highly informative to implicit relationships between UMIs and it might not suffice to find such relationships with custom-built rules especially from a large UMI graph containing a slew of edges. Building on this perspective, we set out to utilize Markov clustering (MCL) [34], a graph-based clustering algorithm, to naturally and spontaneously discover subcomponents with high connectivity from a UMI graph. The MCL-based UMI collapsing process is briefly formulated as follows.

We define *V* as a set of vertices {*v*_1’_*v*_2_,…,*v*_*n*_}representing *n* unique UMIs from raw sequenced reads and *E* as a set of edges {*e*_1’_*e*_2_,…,*e*_*m*_} (*E* ⊂ *V* ×*V*) representing connections between UMIs. Specifically, UMI *v*_*i*_ can connect to UMI *v*_*j*_ as *v*_*i,j*_ if *v*_*i*_ and *v*_*j*_ are *k* edit distance apart. Thus, with a given edit distance *k* we can obtain a graph *G*_*k*_=(V,E)to describe the relationships between UMIs. The adjacency matrix of *G*_*k*_ is denoted as *A*_*k*_ Each element *t*_*i,j*_ of the UMI transition matrix *T*_*k*_ of *G*_*k*_ is generated by applying a standardization operation to each element *a*_*i,j*_ of *A*_*k*_ using the following equation.

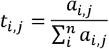

Subsequently, we generate a higher-order transition matrix *D*_*k*_ by taking the power *l* of *T*_*k*_, expressed as

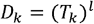

This step is termed the *expansion* of the initial UMI transition matrix *T*_*k*_. Next, we impose an *inflation* operation on the element *d*_*i,j*_ of *D*_*k*_, such that

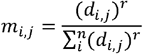

where *r* is an integer. The *inflation* step [35] can enhance the detection of clusters with densely connected UMI nodes and remove weakly connected UMI nodes from clusters. The final transition probability *m*_*i,j*_ in matrix *M*_*k*_ allows clustering of all the raw input UMIs. Based on massive *in silico* permutation tests, we empirically gained that the number of the obtained Markov clusters of UMIs might be large in some cases. For this reason, we further allowed distance *K*_*M*_. Any two Markov clusters are combined if the centre UMIs (*i*.*e*., with the highest the combination between Markov clusters at a UMI cluster level by introducing an edit UMI counts) of the two clusters share a less than *K*_*M*_ edit distance. In the mclUMI tool, this method is dubbed *mcl_ed*. We used *mclumi -m mcl_ed -gt XT -gist XS -ibam (input) -obam (output)* for UMI deduplication throughout this research. Through trials, the expansion parameter *r* is set to 2 and the inflation parameter *l* ranges from 1.4 to 2.

For a bunch of reads, the biggest advantage of mclUMI over established methods (yielding only one UMI deduplication outcome) is the dynamic generation of different count matrices by tunning the *l* (expansion) and *r* (inflation). This property allows mclUMI to search fine-tuned parameters for better deduplication at different situations, e.g., high sequencing errors.

### Post-processing strategies for merging Markov clusters of UMIs

Without considering UMI counts, UMIs are classified into different clusters with MCL based on UMI connectivity. Although the quantity of Markov clusters can be controlled within a certain range by *inflation* and *expansion* as introduced above, there might be association in distances or UMI counts between these clusters. Without considering UMI counts, erroneous UMIs that result from PCR or sequencing errors yet derive from the same ancestor UMI can be corrected by an edit distance to generate a cluster *C*_*a*_ with vertices 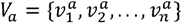. If considering UMI counts, within the cluster, UMI nodes are ideally supposed to evolve from a UMI node 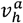 with the highest count and PCR duplicates 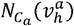 are presumed to be all the neighbouring nodes of 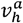, given by

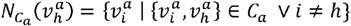

We hypothesised that, at the inter-cluster level, two UMI nodes 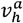 and 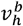 with the highest counts from two clusters *C*_*a*_ and *C*_*b*_ (with vertices 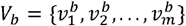) in a small edit distance might be associated in terms of their origin. We further posit that a UMI could be incorrectly synthesised from the UMI of one of the original molecules at the early cycles of PCR amplification and thus gain a high chance to be chosen for amplification at the later cycles of PCR amplification, resulting in an overrepresentation of it in the final sequencing pool. In reality, however, the two UMIs share the same root. Given a set of representative UMI nodes 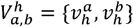, there are *m* + *n* – 1 PCR duplicates, which are identified as the union of their neighbouring nodes 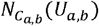 and plus one of themselves. The former part is expressed as

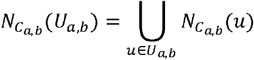

where *C*_*ab*_ is the union of clusters *C*_*a*_ and *C*_*b*_, which is built using the following edit distance.

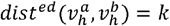

We refer to this edit distance-based strategy as MCL-ed to remove PCR deduplicates through merging of Markov clusters. This parameter is denoted as *mcl_thres*. Inspired by the UMI node merging strategy used in the *Directional* method, we further devise MCL-val, which merges UMI nodes at the Markov cluster (or subcomponents) level. Likewise, PCR duplicates are the neighbouring nodes 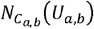 that are considered to be derivatives of 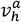 or 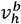 if

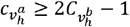

where 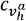 is the count of 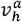 and 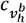 is the count of 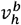. As mentioned above, because two representative nodes can originate from one another at the very early PCR cycles, the following condition is also applicable to the merging of UMIs.

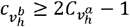

### Representation of UMI sequences

Let *x* be a nucleotide from set *S*^*nt*^ = {*A,T,G,C*}and *S* be a UMI sequence of length *d*. Each nucleotide of *S* is one-hot encoded to be vector *X*^*nt*^ ∈ ℝ ^1×4^

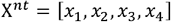

where *x*_*j*_ is given by:

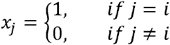

where *i* is the index of a nucleotide in *S*^*nt*^.

Let *X*^*S*^ be the one-hot encoded matrix of *S*. After applied with a flatten operation, a UMI sequence is finally represented by *Y*^*S*^∈ ℝ^1×4*d*^, such that

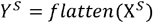

which was fed into Euclidean distance-based clustering algorithms for clustering of UMI nodes that are highly relevant.

### Euclidean distance-based clustering of UMIs

To remove PCR duplicates without building a UMI graph and further compare graph-based deduplication methods, we tested the performance of three Euclidean distance-based clustering algorithms for clustering of UMIs, including density-based spatial clustering of applications with noise (DBSCAN) [36], balanced iterative reducing and clustering using hierarchies (BIRCH) [37], and affinity propagation [38].

#### DBSCAN

Suppose we have *t* unique UMIs *S* = {*S*_1_,*S*_2_,…,*S*_*t*_}observed at a given genomic position and their one-hot encodings are 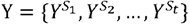. Each UMI has *d* nucleotides. Let *MinPts* be a predefined number of UMIs required for forming a dense region [39]. The Euclidean distance between any two UMIs is computed as follows

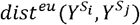

A UMI *S*_*p*_ is said to be a core point if there are at least

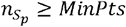

where 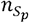 is number of UMIs falling within a region *D*_*ε*_ where any UMI and *S*_*p*_ are within *ε* Euclidean distance, denoted as 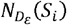

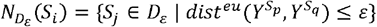

A UMI *S*_*p*_ is said to be a border point if the Euclidean distance between any core point and itself is no greater than *ε* but it has

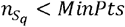

Last, any UMIs that are neither core nor border points are said to be noise points.

DBSCAN starts finding core points, iteratively assigns border points to the dense region of their associated core points, and stops running until all points are labelled a cluster.

#### BIRCH

As BIRCH pertains to complex machinery of clustering data [40], we provide a brief introduction as follow. BIRCH is representative of the hierarchical clusters methods, which utilizes a clustering feature (CF) to describe a cluster, represented as

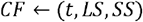

where *t* represents the number of the unique UMIs as given above. *LS* refers to a linear sum operation on the one-hot vectors of UMIs, phrased as

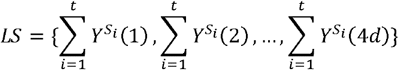

*SS* refers to a squared sum operation on the one-hot vectors of UMIs as follows

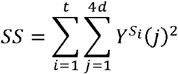

Then, BIRCH builds a CF tree by which CF hierarchical clusters are produced and their cluster centres and scales are updated by computing a few statistics including radii (measured using the Euclidean distance) and diameters.

#### Affinity propagation

Affinity Propagation identifies exemplars of clusters by leveraging three matrices [41], including a similarity matrix *M*^*S*^, a responsibility matrix *M*^*R*^, and an availability matrix *M*^*A*^. Actually, it initially picks a few exemplars as cluster centroids of subsets of UMIs. We take one subset of UMIs *S*_*s*_ = {*S*_1_,*S*_2_,…,*S*_*c*_ | *c* < *t*} ⊂ S as an example to illustrate how to determine an exemplar. We have *M*^*R*^ ∈ ℝ^*c*×*c*^, *M*^*A*^ ∈ ℝ^*c*×*c*^, and *M*^*A*^ ∈ ℝ^*c*×*c*^. 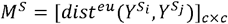 whose elements are the Euclidean distances between UMIs. Elements of *M*^*R*^ and *M*^*A*^ are updated using the following rules.

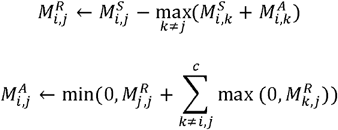

Finally, a UMI at position *i* is chosen to be a cluster exemplar if it satisfies

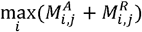

### Evaluation metrics

The method performance for UMI identification is evaluated using the fold change (FC)

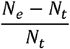

where *N*_*e*_ stands for the estimated number of UMIs and *N*_*t*_ stands for the true number of UMIs. It is used to measure a relative difference.

Alternatively, the performance comparison between methods is evaluated by the difference between fold changes dFC, which is written to be

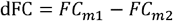

where *m*1 and *m*2 represent two deduplication methods and *FC*_*m*1_ ≤ 0 and *FC*_*m*1_ ≤ 0. dFC > 0 indicates the performance of method *m*1 (or *m*2) better than method *m*2 (or *m*1).

### Settings of simulated reads

We simulated sequencing reads at a single genomic locus to examine the performance of mclUMI. UMI lengths vary from 6 to 18. Reads are amplified from PCR cycles 1-16. Amplification rates are changed from 0.1-1. Sequencing error rates vary in a range [1e-05, 2.5e-05, 5e-05, 7.5e-05, 1e-04, 2.5e-04, 5e-04, 7.5e-04, 1e-03, 2.5e-03, 5e-03, 7.5e-03, 0.01, 0.025, 0.05, 0.075, 0.1, 0.2, 0.3]. Polymerase error rates during PCR amplification are set in a range [1e-05, 2.5e-05, 5e-05, 7.5e-05, 1e-04, 2.5e-04, 5e-04, 7.5e-04, 0.01, 0.05]. Sequencing depths vary in a range [100, 200, 500, 600, 800, 1000, 2000, 3000, 5000]. To make our experiment controllable, we generated sequencing data using one of the above parameter settings each time while keeping the rest of parameters fixed. Based on our empirical analysis and in line with the *UMI-tools* work, the fixed parameters of PCR cycles, PCR errors, sequencing errors, UMI lengths, and amplification rates were set to 8, 1e-05, 1e-03, 10, and 0.85, respectively. To eliminate random errors and biases, we carried out 10 permutation experiments over which the final fold change is averaged.

### Formulation of read simulation

The total number of reads 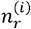 at PCR cycle *i* consists of two sources of reads, i.e., reads that are and are not PCR amplified. An initial number of reads is set to 50. The estimate of amplified reads follows a Galton-Watson branching (GWB) process [42,43], such that

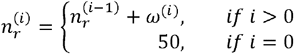

where *ω*^(*i*)^ is the total number of reads resulting from PCR amplification and can be calculated from a binomial distribution

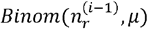

where *μ* is the amplification efficiency.

Likewise, we use the same way to simulate PCR errors. The total number of errors 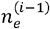 at PCR cycle *i* is calculated as

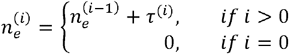

where *τ*^(*i*)^ is the number of errors synthesised at PCR cycle *i* and is drawn from a negative binomial distribution [44]

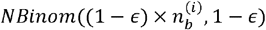

where *ϵ* is the PCR error rate and 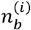 is the number of bases of amplified reads at PCR cycle *i*.

The number of sequencing errors *n*_*e*_ is estimated from a negative binomial distribution

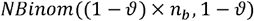

where *∂* is the sequencing rate and *n*_*s*_ is the total number of bases after PCR amplification.

### Simulation of single-cell count matrices

We employed SPsimSeq [29] to generate an expression matrix of genes in single cells simulated based on the SingleCellExperiment data [45]. We simulated a single-cell count matrix with 10 genes and 10 cells to create an initial pool of 1828 unique molecules attached with cell barcodes and UMIs. The length of simulated UMIs is set to 10 in line with the protocol of10X Genomics V2 chemistry.

To supply information about cell types, we then used DeepConvCVAE trained by a variational autoencoder on the PBMC68k dataset [30] (https://github.com/10XGenomics/single-cell-3prime-paper/tree/master/pbmc68k_analysis) and generated a count matrix of 1100 cells (100 cells per each of 11 cell types) and 20,736 genes. The DeepConvCVAE framework and the trained model are currently housed in UMIche (https://2003100127.github.io/umiche).

### Identification of highly expressed genes

We identified the top *k* highly expressed genes by

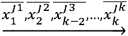

where 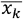 represents the identifier of a gene with the *k* th highest average gene expression value at column *jk* (*jk* ∈ 1,2,…,*n*)of a UMI count matrix with *m* cells and *n* genes.

## Code and data availability

The mclUMI software is publicly available at https://github.com/cribbslab/mclumi.

## Author contribution

J.S. conceived this research. J.S. developed the algorithm, implemented mclUMI, and wrote the manuscript. J.S. conducted data analysis with contributions from S.L., S.C., and A.P.C. J.S. and A.P.C. acquired funding. All authors have given approval to the final version of the manuscript.

## Acknowledgement

This work was financially supported by the Medical Research Council (MRC) career development fellowship (MR/V010182/1).

