## Supplementary Information for "mclUMI: Markov clustering of unique molecular identifiers enables dynamic removal of PCR duplicates"

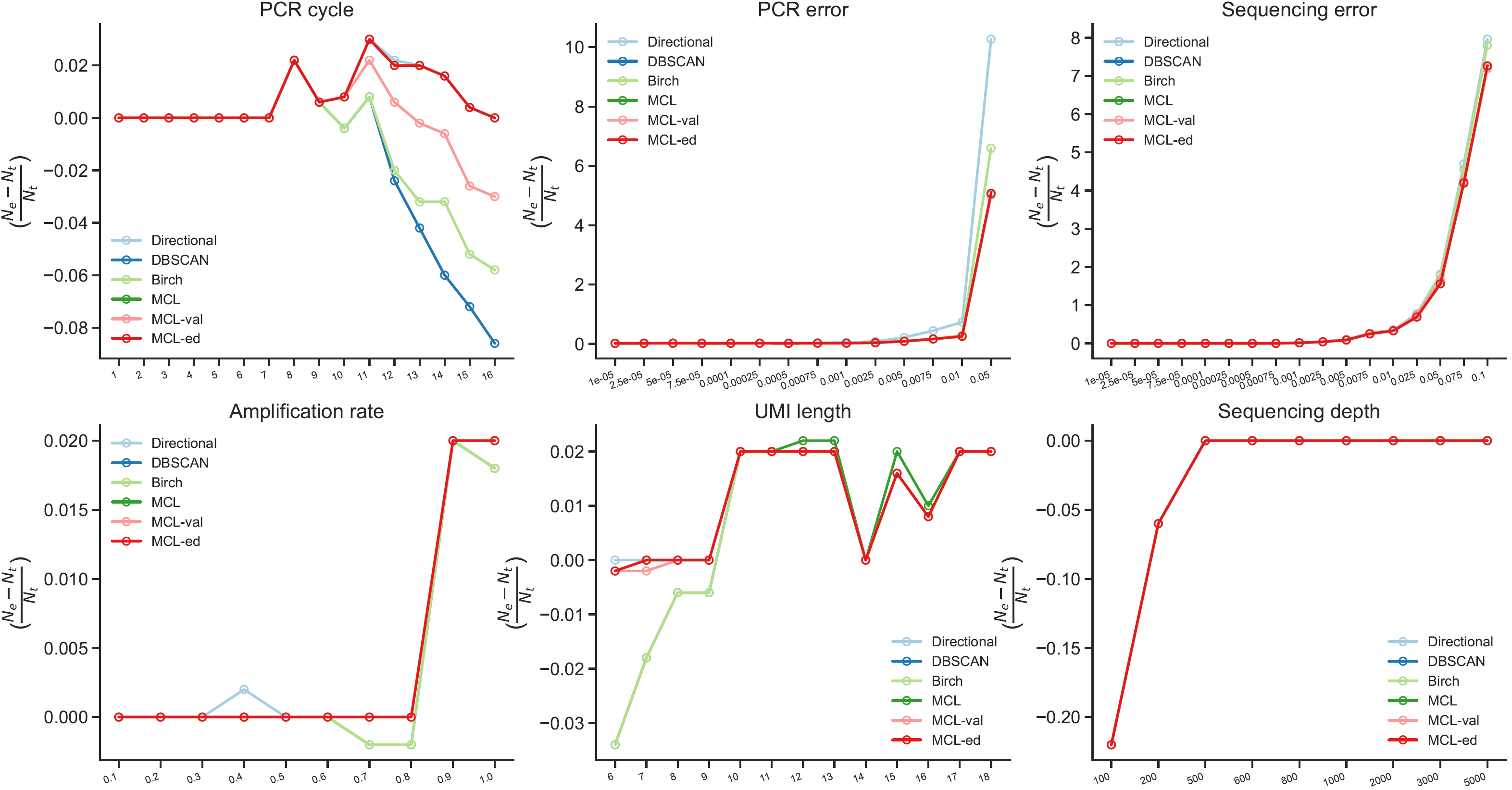

**Supplementary Figure 1.** Fold change between the actual number of UMIs and the number of UMIs deduplicated by Directional, MCL, MCL-ed, MCL-val, DBSCAN, and Birch across multiple PCR cycles, PCR error rates, sequencing error rates, amplification rates, UMI lengths, and sequencing depths, respectively.

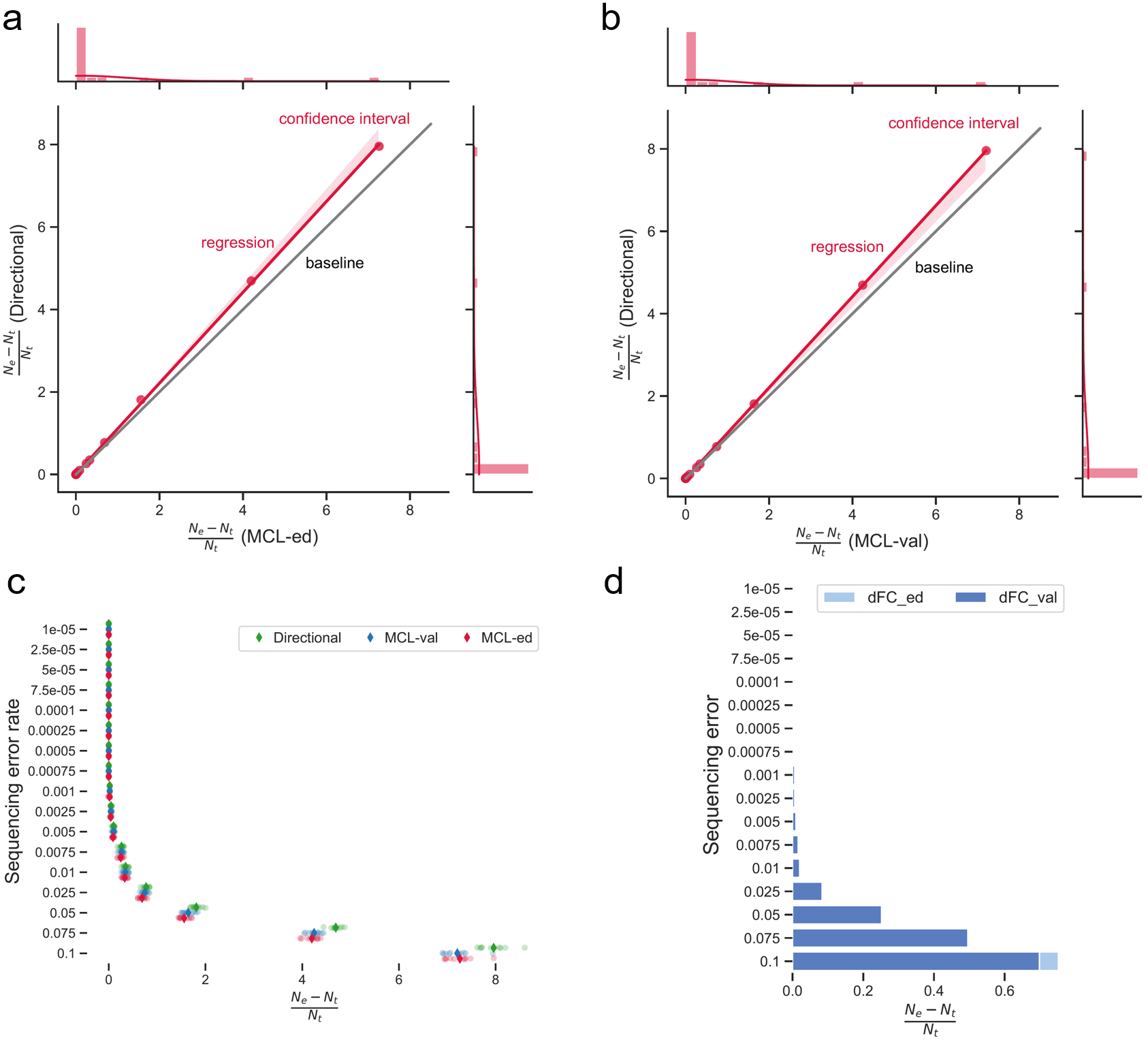

**Supplementary Figure 2.** Comparison of deduplication results per permutation test between mclUMI (MCL-val and MCL-ed) and the UMI-tools *Directional* method across multiple sequencing error rates. **a** and **b**, Directional FCs plotted against mcl_ed FCs (**a**) and mcl_val FCs (**b**). Lines with confidence intervals in red are plotted by fitting the paired FCs using regression functions. **c**, Strip plots display the averaged FC values in diamonds and the FCs per permutation test in circles. **d**, Method comparison using the difference of fold changes (dFCs). dFC_ed represents the fold changes between using the *Directional* method and MCL-ed and dFC_ val represents the fold changes between using the *Directional* method and MCL-val.

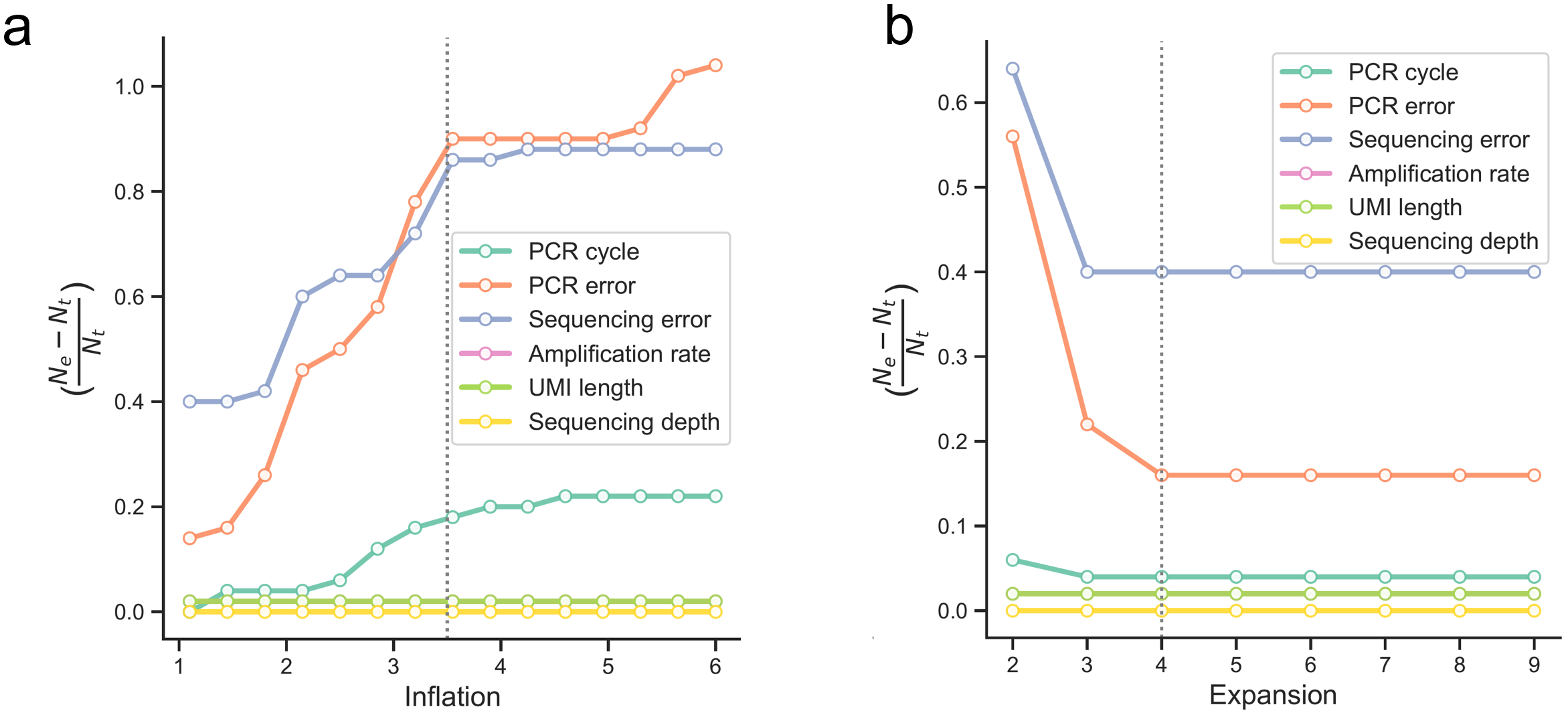

**Supplementary Figure 3.** Fine-tuned parameters of the MCL algorithm. **a** and **b**, Fold change between estimated and actual deduplicated counts with respect to the inflation (**a**) and expansion (**b**) parameters of the MCL algorithm.
